# Critical role of H3K27 methylation in cell fate determination of two cell lineages in male gametophyte

**DOI:** 10.1101/2021.11.15.468715

**Authors:** Xiaorong Huang, Meng-Xiang Sun

## Abstract

During angiosperm male gametogenesis, microspores divide to produce a vegetative cell (VC) and a male germline (MG), each with a distinct cell fate. How the MG cell/VC fate is determined remains largely unknown. Here, we report that H3K27me3 is essential for VC fate commitment and H3K27me3 erasure contributes to the MG cell fate initiation in male gametophyte of *Arabidopsis*. The VC-targeted H3K27me3 erasure disturbed the VC development and resulted in the VC fate shifting towards a gamete destination, which suggests that MG cells require H3K27me3 erasure for triggering the gamete cell fate. Multi-omics and cytologic analysis confirmed the occurrence of extensive cell identity transition due to H3K27me3 erasure. Therefore, we experimentally confirm that the MG cell/VC fate is epigenetically regulated. The H3K27 methylation plays a critical role in the guidance of MG cell/VC fate determination for pollen fertility in *Arabidopsis*. Our work also provides new evidences for two previous hypotheses that the germline cell fate is specified by the differential distribution of yet unknown determinant, and VC maintains the microspore’s default program, namely the H3K27me3 setting, whereas MG requires reprogramming.

## Introduction

During sexual development in angiosperms, the factors that determine the cell fates of male germline (MG) cells and vegetative cells (VCs) remain largely unknown. Previous reports have shown that the asymmetric division of angiosperm microspores is decisive for MG cell formation and specification (Twell et al., 1998). MG cell fate is specified by unknown factors confined in the smaller daughter cell following asymmetric division of the microspore (Twell et al., 2002; Hackenberg and Twell, 2019). Mutation of *RETINOBLASTOMA RELATED* (*RBR*) in microspores causes hyperproliferation and prevents the establishment of cell identity in VCs. The VCs of *rbr* mutants exhibit non-specific asymmetric division, producing an additional cell with a VC fate, MG cell fate, or mixed cell fate (Chen et al., 2009). Once the MG cell type is determined, it produces male gametes that follow a developmental program that is unaffected by the cell cycle or the VC-supplied microenvironment (Chen et al., 2008; Huang et al., 2018). Notably, the MG cell-specific conserved pathway mediated by R2R3 Myb-domain protein DUO POLLEN1 (DUO1) is essential for triggering MG cell specification (Rotman et al., 2005; Berger and Twell, 2011; Borg et al., 2011; Higo et al., 2018). These studies suggest that the cell fate dichotomy of VCs and MG cells is triggered after asymmetric cell division of the microspore. However, how germline fate is suppressed in VCs but established in MG cells is currently unknown; similarly, the determining factors that promote *DUO1* expression in MG cells have yet to be identified. Recently, to answer the question, epigenetic regulation has drawn a great attention.

DNA methylation is globally erased in the paternal chromatin of mammals; by contrast, DNA is not demethylated at least in CG and CHG context in the *Arabidopsis* MG cells (Calarco et al., 2012; Borg and Berger, 2015). In further contrast to mammals, the sperm chromatin of which undergoes widespread removal of histone modifications as the histones are replaced with protamines (Reik W et al., 2001; Sassone-Corsi, 2002), H3.1 histones are replaced by the MG cell-specific histone variant H3.10 (HTR10) in the sperm chromatin of flowering plants (Ingouff et al., 2010). H3.10 is unable to receive H3K27me3 modifications, and its even incorporation in sperm chromatin coupled to the activity of the H3K27me3 demethylases RELATIVE OF ELF 6 (REF6), EARLY FLOWERING 6 (ELF6) and JUMONJI 13 (JMJ13) leads to a global loss of H3K27me3 (Borg et al., 2020), suggesting H3K27me3 erasure may play a critical role in MG cell fate determination. However, the sperm cells (SCs) were generated in both *elf6 ref6 jmj13 htr10* quadruple mutant and the SC-ectopic *SWINGER* (*SWN*, methyltransferase subunit of Polycomb repressive complex 2 (PRC2)) expression lines (Borg et al., 2020). Therefore, the role of H3K27 methylation in cell fate determination of two cell lineages in male gametophyte remains to be demonstrated experimentally. In contrast to MG cells, the chromatin of VCs undergoes global loss of the centromeric H3 variant CENH3 and the heterochromatic mark H3K9me2, both of which persist in MG cells (Schoft et al., 2009). *Arabidopsis SET DOMAIN GROUP2* (*SDG2*) mutation disrupts H3K4me3 deposition, which leads to ectopic H3K9me2 speckle formation and chromatin condensation in VCs. However, this does not alter VC or SC fate establishment (Pinon et al., 2017), suggesting that both H3K4me3 and H3K9me2 act primarily on chromatin remodeling, as opposed to cell fate determination during male gametogenesis. To sum up, whether the histone posttranslational modifications (PTMs) guide cell fate determination in male gametophytes still remains unclear. Herein, we experimentally confirmed that the H3K27me3 is essential for maintaining VC fate and H3K27me3 erasure resulted in VC fate shifting to the MG cell fate, suggesting its critical role in the guidance of MG cell/VC fate determination for pollen fertility in *Arabidopsis*.

## Results

### H3K27me3 is essential for VC functional specification

In wild-type (WT) mature pollen, the strong immunofluorescence signals exhibiting H3K27me3 were detected in VC nuclei (VN) but not in SC nuclei (SN), indicating that distinct H3K27 methylation modification between VC and SC (Figure 1A). To investigate the role of H3K27 methylation in cell fate determination during male gametophyte development, we first observed the male gametophyte development in H3K27 methyltransferases mutants. According to previous reports the fertile double homozygous mutant of *SWN* and *CURLY LEAF* (*CLF*), methyltransferase subunits of PRC2, could not survive (Lu et al., 2011). The analysis of expression pattern showed that *SWN* and *CLF* were expressed in the microspore, but only *SWN* was sequentially expressed in VCs after asymmetric division of the microspore (Borg et al., 2020). *SWN* may play a major role in H3K27me3 maintenance in VCs, and thus, *swn-21/-clf-29/+* was used legitimately in our experiments. Although the germination rate was reduced along with the decrease of H3K27me3 level in VCs, there was no obvious morphological abnormality in the *swn-21/-clf-29/+* male gametophytes (Supplemental Figure S1). Since H3K27me3 was not completely erased in the *swn-21/-clf-29/+* VCs (Supplemental Figure S1, C and D), to better explain the role of H3K27me3 in the fate determination of distinct cell lineages in male gametophytes it is necessary to create more suitable materials.

**Figure 1.**
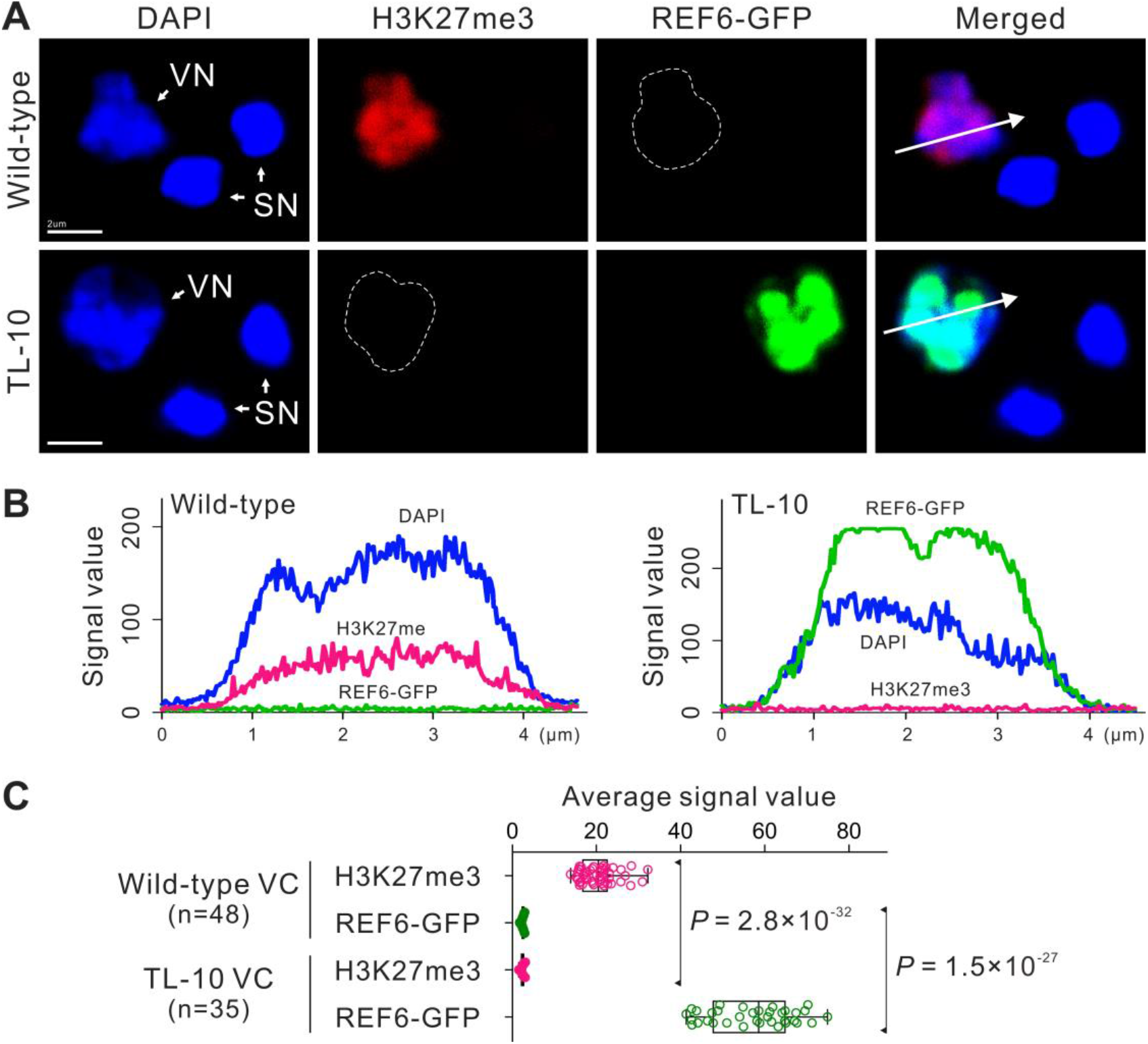
VC-targeted H3K27me3 erasure in male gametophyte. **A**, *REF6* was expressed under the control of *Lat52* promoter to purposefully erase H3K27me3 in VCs. The transgenic lines were named as REF6-Transferred Line (REF6-TL, hereafter TL). Whole-mount H3K27me3 antibody labeling and DAPI staining of mature pollen in wild-type (WT) (top) and strong TL-10 line (bottom). Dotted lines indicate vegetative nucleus (VN). SN, sperm cell nucleus. **B**, Fluorescence signal intensity of WT (left) and TL-10 (right) VN are displayed along the direction of the long white arrow in (**A**). **C**, Statistical analysis of the average signal intensity of REF6-GFP and H3K27me3 in VN from WT and TL-10. Scale bars, 2μm (**A**).

Known that dynamic equilibrium of histone lysine methylation is maintained by the histone lysine methylases and demethylases (Liu et al, 2010), we generated a transgenic *Arabidopsis* line that expresses *REF6*, the first H3K27 demethylase identified in plants (Lu et al., 2011; Cui et al, 2016). *REF6* was placed under control of the *Lat52* promoter to purposefully erase H3K27me3 in VCs. The transgenic line was named as REF6-Transferred Line (REF6-TL, hereafter TL). The H3K27me3-specific fluorescent signal was barely detectable in VNs of strong TL-10 line, indicating almost disappearance of H3K27me3 (Figure 1). Homozygous could not be obtained in the strong TL lines (TL-09/+, −10/+, −18/+ and −30/+) (Supplemental Figures S2, A-D and S3). A typical function of VCs is to germinate and form a pollen tube to deliver SCs to the embryo sac for double fertilization; we therefore examined this function in various TL lines. No obvious differences in viability or morphology were observed in the strong TL line pollen compare with the control; however, the mature pollen from the strong TL lines showed abnormal vesicle organization and failed to germinate (Figure 2, A-J; Supplemental Figures S2, D-I and S3, A and B). In addition, similar to *swn-21/-clf-29/+*, the morphology of pollen was normal, but the pleiotropic phenotypes of pollen germination were observed in various weak homozygous TL lines (TL-13/-, −21/- and −25/-). In the TL-13 line, ∼60% of pollen lost the ability to germinate, and ∼37% of pollen tubes burst immediately after germination; only ∼3% tubes remained intact but elongated slowly compared with control. Thus, the TL-13 line produced few seeds. Similar defects were exhibited by the other weak TL lines (Supplemental Figures S2, B and J and S3, C-E and K-M). Interestingly, H3K27me3 removal disrupted the formation of the male germ unit (MGU), which is a typical feature of a functional VC (Figure 2, K-M). These results suggest that H3K27me3 is essential for VC developmental stability and functional specification.

**Figure 2.**
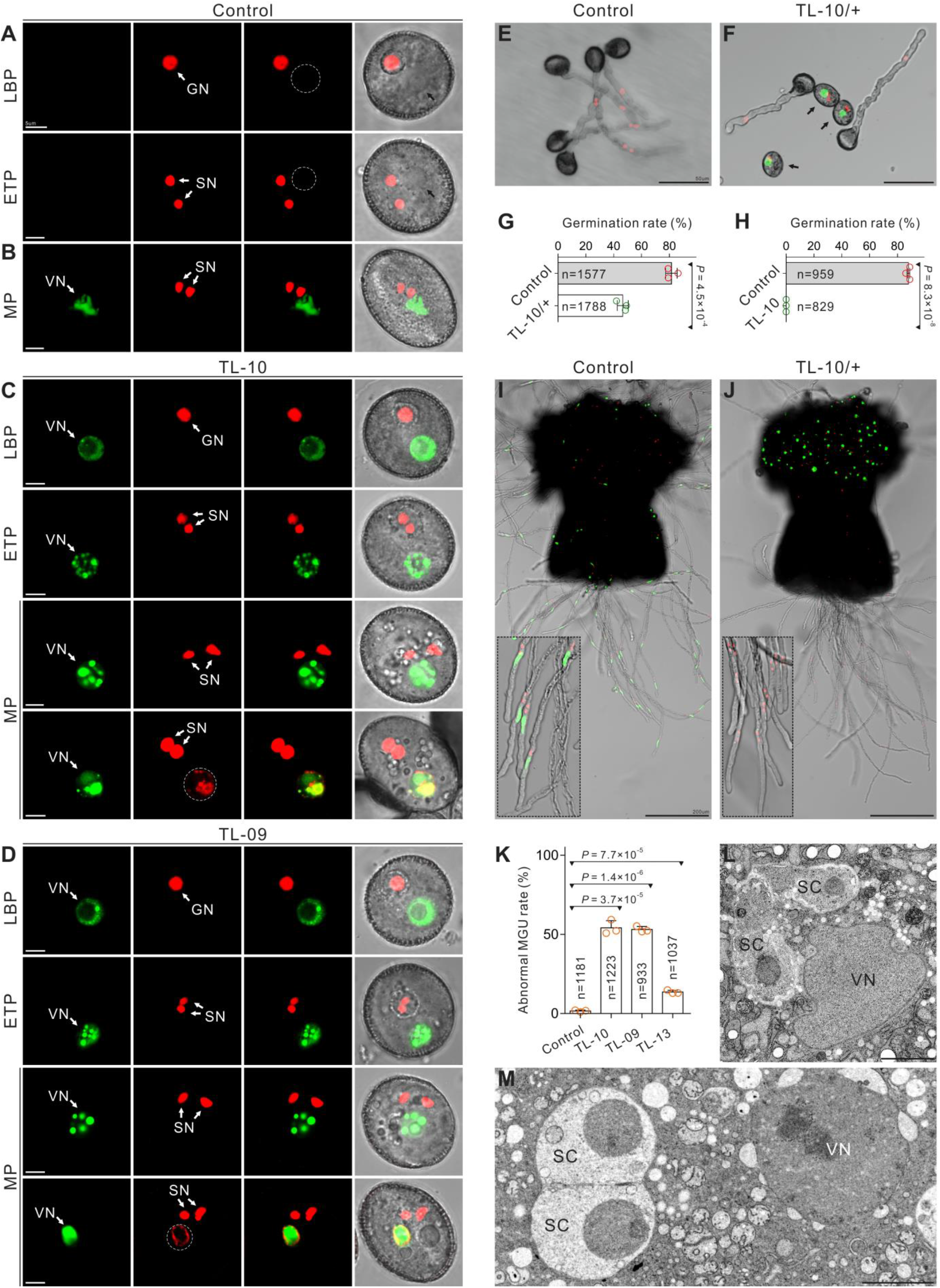
H3K27me3 is essential for VC identity and functional specification. **A, B**, Normal morphology in control VCs from the *pHTR10:HTR10-RFP* line (**A**) and *pACTIN11:H2B-GFP*; *pHTR10:HTR10-RFP* double-labeled line (**B**). Dotted lines and black arrows indicate vegetative cell nucleus (VN). **C, D**, The morphology of the strong TL VCs at different development stages. SC formation and VC viability were unaffected, but the abnormal vesicle organization and ectopic HTR10-RFP signal were detected in the mature pollen from TL-10 (**C**) and TL-09 (**D**) lines. Dotted lines indicate the ectopic HTR10-RFP signal in VN. **E, F**, *In vitro* pollen germination assay of *pHTR10:HTR10-RFP* line (**E**) and TL-10/+ carrying the HTR10-RFP marker (**F**). Arrows indicate the pollen unable to germinate. **G, H**, Comparison of the *in vitro* pollen germination rate between control and TL-10/+. Results are from three independent experiments. **I, J**, *In vivo*-*in vitro* pollen germination assay following pollination of WT pistils with control pollen (*pACTIN11:H2B-GFP*; *pHTR10:HTR10-RFP* double-labeled line (**I**)) and TL-10/+ pollen (**J**), respectively. Dotted box in (**I**) displays the male germ unit (MGU) in pollen tubes. Dotted box in (**J**) indicates that TL-10 pollen failed to form tubes and all the observed tubes are from normal pollen. **K**, Statistical analysis of the abnormal MGU construction in control and TL lines. Results are from three independent experiments. Statistical significance is determined by Student’s t-test (P < 0.001, ***) and the error bars show standard deviation (SD). Transmission electron microscopy (TEM) of WT (**L**) with normal MGU and TL-10 (**M**) pollen with abnormal MGU construction. Scale bars, 5μm (**A-D**), 50μm (**E, F**), 200μm (**I, J**), 2μm (**L, M**). GN, generative cell nucleus; SN, sperm cell nucleus; SC, sperm cell; VN, vegetative cell nucleus; LBP, late bicellular pollen; ETP, early tricellular pollen; MP, mature pollen.

### VC-targeted H3K27me3 erasure results in the VC cell fate shifting towards MG

Exhilaratingly, we found the aberrant phenotypes of the strong TL pollen grains were similar to the previously reported VC-ectopic *DUO1* expression line (*pLat52:DUO1*) (Brownfield et al., 2009), Thus, the VC-targeted H3K27me3 erasure likely changed the attributes of the strong TL VCs and triggered a cell fate transition to the MG. To test the TL VC cell identity, we examined the MG cell-specific markers in the strong TL pollen. As expected, the HTR10-RFP marker localized in ∼11% VN of TL-09, −10, −18 and −30 pollen; this was not observed in WT VN (Figure 2, A-D and 3, A and D; Supplemental Figure S2, H and I). DUO1 localization was examined using DUO1-RFP. Ectopic DUO1 signals were detected in ∼16% VN of TL-09, −10, −18 and −30 pollen (Figure 3, B and D). These findings suggest that the targeted removal of H3K27me3 in VCs induced a cell fate shift in the VC to MG cell. In plants, the H3K9me2 modification is enriched in hypermethylated heterochromatin. H3K9me2 and methylated CHG interact in a self-enforcing positive feedback loop that acts on constitutive heterochromatin and may contribute to extreme chromatin condensation in the SN (Borg and Berger, 2015; Schoft et al., 2009; Jackson et al., 2002; Johnson et al., 2002). Here, we found that the H3K9me2 marker was present only in the SN of WT pollen, whereas H3K9me2 was detected in both SN and VN in 9.90% (± 0.80) of TL-10 pollen (Figure 3, C and D); the chromatin H3K9me2 and H3K27me3 status of TL-10 VN was similar to that of SN, suggesting a cell identity transition. Therefore, our results suggest that the MG cell fate determination is epigenetically regulated, and H3K27me3 erasure is one of the important factors to induce the MG cell fate.

**Figure 3.**
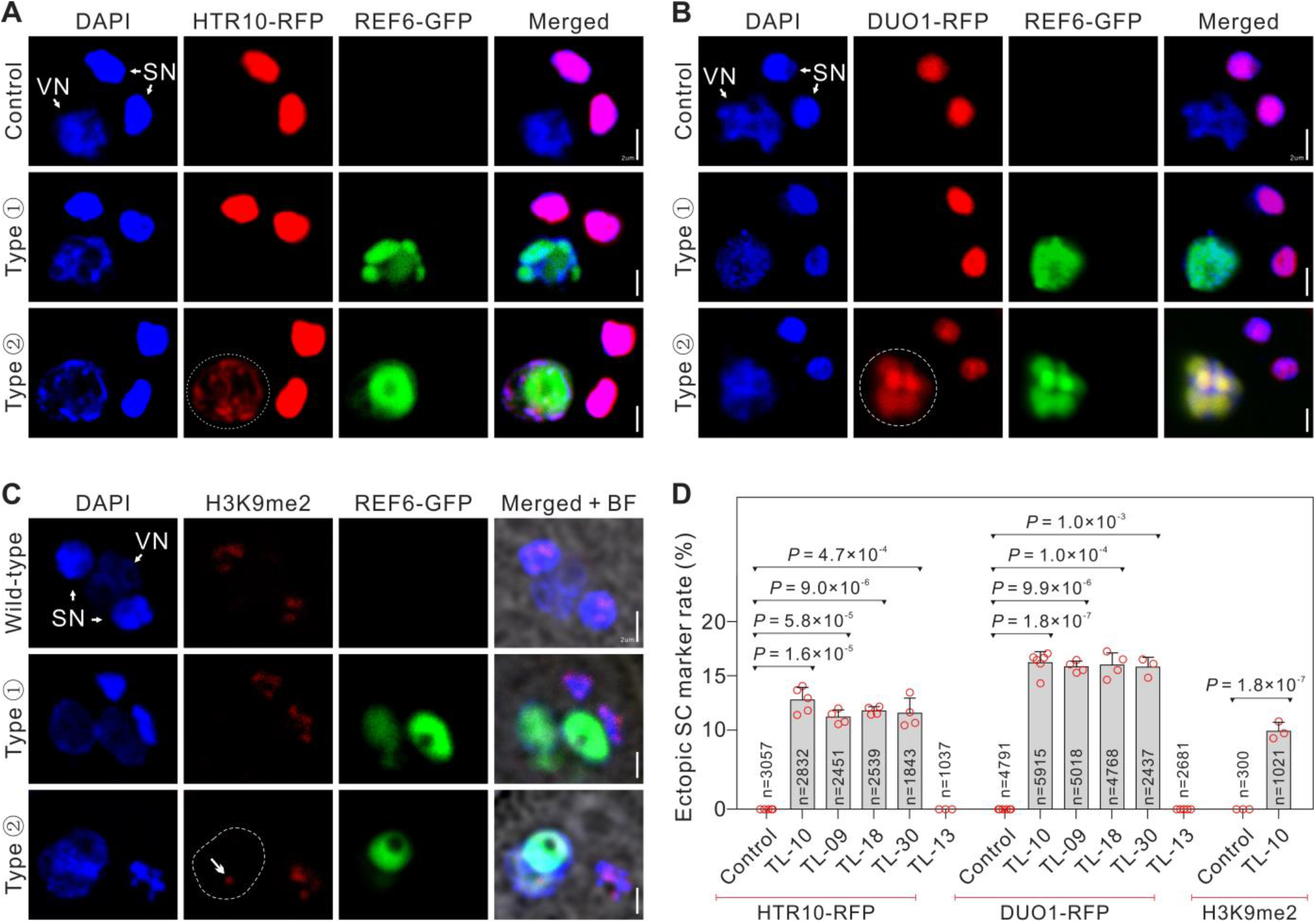
VC-targeted H3K27me3 erasure triggers the VC-SC fate transition. **A-C**, The MG cell-specific markers in control and TL-10 pollen. Fluorescent images of the male germ unit (MGU) stained with DAPI, also indicated by pLat52:REF6-GFP and pHTR10:HTR10-RFP (**A**), pDUO1:DUO1-RFP (**B**) and H3K9me2 signals (**C**), in control and TL-10. Type ➀ and type ➁ indicate TL-10 VN without or with MG cell-specific marker signals, respectively. Dotted lines indicate the ectopic HTR10-RFP (**A**), DUO1-RFP (**B**) signals, and arrow indicates the ectopic H3K9me2 signals in VN (dotted line) (**C**). **D**, Statistical analysis of the ectopic MG cell-specific marker signal in VN. Statistical significance is determined by Student’s t-test (P < 0.001, ***) and the error bars show standard deviation (SD). Scale bars, 2μm (**A-C**).

### H3K27me3 erasure leads to chromatin remodeling in TL VCs

To further investigate the impact of the H3K27me3 erasure on TL VC cell fate, the pure VNs and SNs were isolated from the broken pollen mixture via fluorescence activated nuclei sorting (FANS) for H3K27me3 cleavage under targets and tagmentation (CUT&Tag) and assay for transposase accessible chromatin (ATAC)-seq analyses. In total, six CUT&Tag and eight ATAC-seq libraries were constructed and sequenced (Supplemental Figures S4 and S5). Principal component analysis (PCA) of the CUT&Tag and ATAC-seq datasets showed that profiles of the same cell lineage were consistent and well separated from each other (Figure 4, A and B). H3K27me3 is generally enriched in chromatin areas containing silenced genes. The CUT&Tag results revealed 6950 H3K27me3 peaks in WT VCs, by contrast, WT SCs generated 694 peaks which was similar to the previously reported H3K27me3 ChIP-seq data of WT SCs (Borg et al., 2020); TL VCs generated 141 peaks, indicating that H3K27me3 modifications were effectively erased in TL VCs, and that the H3K27me3 state of TL VCs closely resembled that of WT SCs (Figure 4, C-F; Supplemental Figure S4, E-G). For the ATAC-seq analysis, ∼16,000 peaks were identified in WT and TL SCs, and ∼21,000 peaks were detected in WT and TL VCs; the data quality was therefore sufficient for the following analyses.

**Figure 4.**
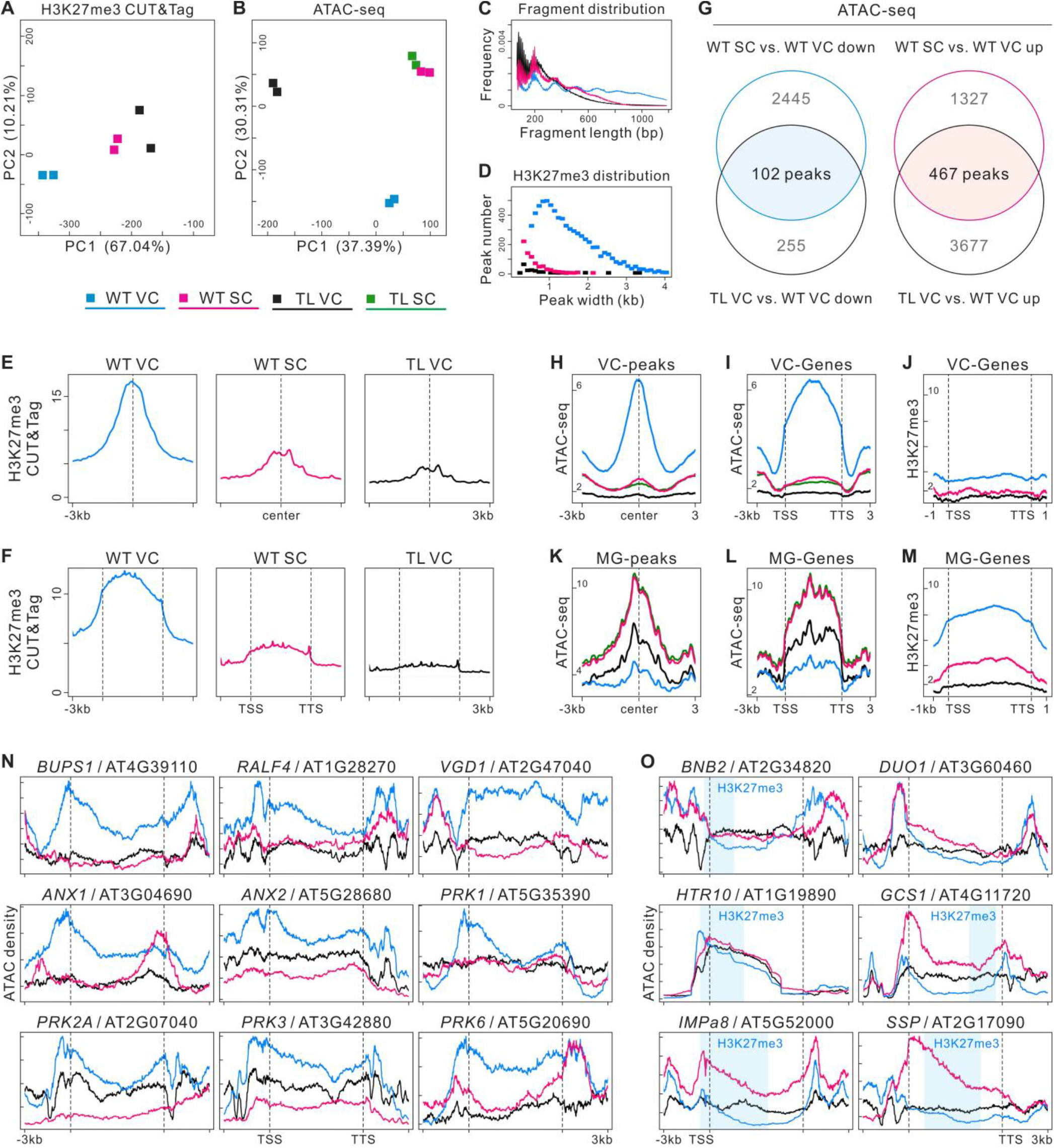
H3K27me3 erasure leads to chromatin remodeling in TL VC. **A, B**, Principal component analysis (PCA) of H3K27me3 CUT&Tag (**A**) and ATAC-seq data (**B**), illustrating high reproducibility of replicates and the relationship among cell lineages. TL-10/+ was used in experiments. **C**, Sample fragment length distribution of H3K27me3 CUT&Tag library reads. **D**, Box plot of the size distribution (in kb) of WT VC, TL VC, and WT SC H3K27me3 domains. The peaks numbers were 6950 in WT VCs, 694 in WT SCs, and 141 in TL VCs. **E, F**, CUT&Tag profiles over H3K27me3 domains of WT VC in three cell lineages respectively (**E**) and H3K27me3 signals over peak-annotated genes (**F**) suggest that H3K27me3 is maintained in WT VCs but lost in WT SCs. Similar to WT SCs, H3K27me3 levels are low in TL VCs. CUT&Tag data were generated from two biological replicates. **G**, Overlap analysis of down-regulated (left) and up-regulated (right) ATAC-seq peaks between WT SC vs. WT VC (WT SC compared with WT VC) and TL VC vs. WT VC. The overlapped regions indicate VC-required peaks (VC-peaks) and MG-required peaks (MG-peaks). **H, I**, ATAC-seq profiles over VC-peaks (**H**) and chromatin accessibility signals over peak-annotated genes (VC-genes) (**I**) in four cell lineages. **J**, H3K27me3 signals over the VC-genes annotated by VC-peaks. **K, L**, ATAC-seq profiles over MG-peaks (**K**) and chromatin accessibility signals over peak-annotated genes (MG-genes) (**L**) in four cell lineages. **M**, H3K27me3 signals over the MG-genes annotated by MG-peaks in promoter transcriptional start sites (promoter-TSS), gene bodies (exons and introns), and transcription termination sites (TTS), indicating that these genes are repressed by H3K27me3 in WT VCs and activated by H3K27me3 erasure in WT SCs and TL VCs. **N, O**, Genome browser views of merged ATAC-seq datasets from WT VC, TL VC, and WT SC; chromatin accessibility was low in VC-related genes (**N**) and high in MG cell-related genes (**O**) of TL VCs. The chromatin accessibility of MG cell-related genes changed near H3K27me3 domains (light blue areas) along gene bodies.

We then aimed to determine whether chromatin regions were altered during MG cell specification and how this is related to the targeted H3K27me3 erasure. Pairwise comparisons of the ATAC-seq datasets revealed 1794 differentially accessible regions (DARs) with increased openness (up) and 2547 DARs with decreased openness (down) in WT SCs compared with WT VCs (WT SC vs. WT VC). In TL VCs compared with WT VCs, 4195 DARs displayed significantly higher accessibility and 357 DARs decreased accessibility (Supplemental Figure S5, F and G). Gene ontology analysis revealed that the genes displaying peaks “WT SC vs. WT VC down” or “TL VC vs. WT VC down” were related to pollen tube development, such as pollen tube growth, regulation of pH, and receptor kinase signaling. Furthermore, the genes displaying peaks “WT SC vs. WT VC up” or “TL VC vs. WT VC up” were related to SC differentiation, as well as cell–cell signaling and cell fate commitment (Supplemental Figure S5H); this suggests an accessibility modification of specific chromatin region for the cell fate transition in TL VCs.

Next, we combined the ATAC-seq data from the WT SC vs. WT VC and TL VC vs. WT VC comparisons, which yielded 102 VC-required peaks (VC-peaks) and 467 MG-required peaks (MG-peaks) related to cell lineage specificity in male gametophytes (Figure 4, G, H and K). We found that the annotated genes associated with VC-peaks were less accessible in WT SCs and TL VCs than in WT VCs. On the other hand, the annotated genes associated with MG-peaks tended to be more open in WT SCs and TL VCs than in WT VCs (Figure 4, I and L; Supplemental Figure S6F). In WT VCs, the annotated genes of the VC-peaks harbored fewer H3K27me3 modifications, and the annotated genes of the MG-peaks were broadly suppressed by H3K27me3, especially in gene body and promoter-transcription start sites (Figure 4, J and M; Supplemental Figure S6E).

Previous reports identified essential VC- and MG cell-related genes, which we used as markers in the following work. The well-established VC-preponderant or -specific genes *ANX1*/*2, BUPS1*/*2, RALF4, PRK1/2A/3/6* and *VGD1*, which are involved in pollen tube growth and guidance (Jiang et al., 2005; Boisson-Dernier et al., 2009; Takeuchi and Higashiyama, 2016; Ge et al., 2017), were more accessible in WT VCs than in WT SCs, and less accessible in TL VCs (Figure 4N). Conversely, the MG cell-specific genes, which include *BNB2, DUO1, HTR10, GCS1, SSP*, and *IMPa8* (Bayer et al., 2009; Borg et al., 2011; Yamaoka et al., 2018), displayed a less accessible chromatin status in WT VCs than in WT SCs, whereas their accessibility was higher in TL VCs (Figure 4O). In WT VCs, the VC-preponderant or -specific genes were not marked with H3K27me3, but the MG cell-specific genes (with the exception of *DUO1*) were marked with H3K27me3 in the closed chromatin region (Figure 4, N and O; Supplemental Figure S4H), suggesting that H3K27me3 participates in VC identity maintenance, and that its erasure may trigger MG cell differentiation. Although *DUO1* itself was not modified by H3K27me3, its potential upstream regulator BNB1/2 (Yamaoka et al., 2018) was marked by H3K27me3 (Supplemental Figure S4H). Thus, H3K27me3 erasure may activate, directly or indirectly, the transcriptional pathways of MG cell specification via a pleiotropic mechanism. VC-targeted H3K27me3 erasure therefore led to VC-related chromatin closure and MG cell-related chromatin opening in TL VCs.

### H3K27me3 erasure induces transcriptome reconstruction in TL VCs

To further assess which genes were regulated respectively in VC and MG cell fate determination, VN and SN were isolated for RNA-seq, and twelve transcriptome libraries were constructed and sequenced (Supplemental Figure S6A). In total, ∼20,000 genes were detected in the transcriptomes of WT/TL SCs and WT VCs and ∼27,000 in TL VCs. PCA of the transcriptomic datasets showed that the profiles of the same cell lineage were consistent, and PC1 and PC2 were sufficient to separate the samples into four groups. Although TL VCs and WT VCs were clearly separated, TL SCs and WT SCs were clustered more closely (Figure 5, A-C). Moreover, correlation matrices showed that TL VCs were more similar to WT SCs compared with WT VCs at the transcriptional level (Figure 5B), suggesting a transcriptome transition from VCs to SCs due to the targeted H3K27me3 erasure. These results suggest that H3K27me3 erasure altered the nature of TL VCs but did not significantly affect the identity of TL SCs; this was further supported by unsupervised hierarchical clustering (UHC) analysis of the transcriptomic datasets (Supplemental Figure S6D). Since the *Lat52* promoter weakly drives the *REF6* expression at late microspore stage, six transcriptome libraries from the late microspore (LMS) of both WT and TL-10 were sequenced to assess possible effect of the week *REF6* expression on succedent cell lineages specialization (Supplemental Figures S2, D-G and S6B). The correlation matrices and UHC analysis showed that the transcriptomic datasets of WT LMSs and TL LMSs were clustered together (Supplemental Figure S6, C and D), indicating the weak *REF6* expression had no significant effect yet on TL-10 LMS development, which was consistent with aforesaid phenotypic analysis.

**Figure 5.**
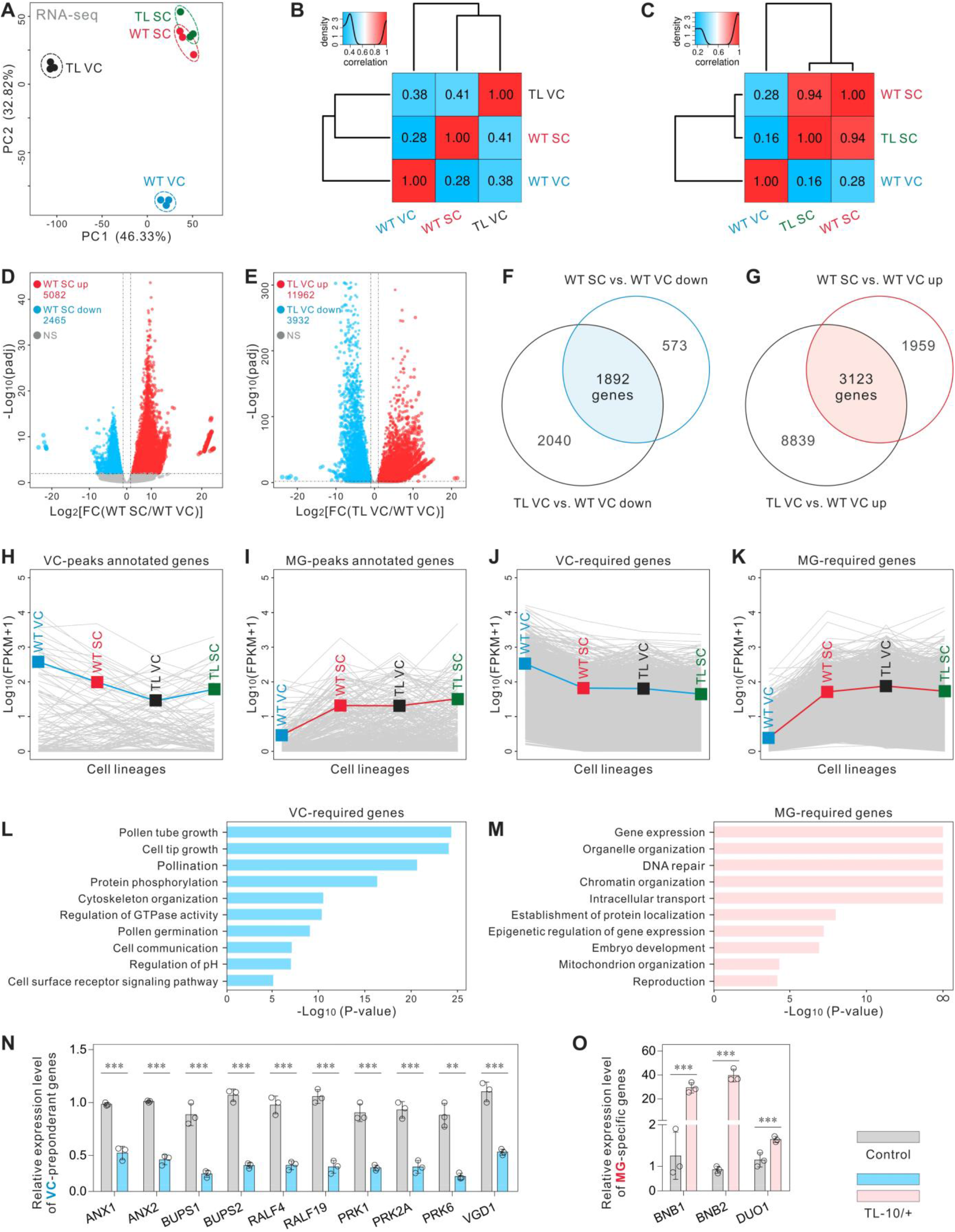
H3K27me3 erasure induces transcriptome reconstruction in TL VCs. **A**, PCA of WT VC, WT SC, TL VC, and TL SC transcriptomes, illustrating the high reproducibility of replicates and the relationship among the four cell lineages. TL-10/+ was used in experiments. **B**, Pearson correlation matrix of the WT VC, WT SC, and TL VC transcriptomes shows that TL VC is more similar to WT SC, compared with WT VC. **C**, Pearson correlation matrix of the WT VC, WT SC, and TL SC transcriptomes shows that there is no significant difference between TL SC and WT SC. **D, E**, Scatterplot showing the numbers of up- and down-regulated genes in WT SC vs. WT VC (**D**) and TL VC vs. WT VC (**E**). **F, G**, Overlap analysis of the down- (**F**) and up-regulated (**G**) genes between WT SC vs. WT VC and TL VC vs. WT VC. The overlapped regions indicate VC-required genes (**F**) and MG-required genes (**G**), respectively. **H, I**, Expression profiles of VC-peak annotated genes (**H**) and MG-peak annotated genes (**I**) in four cell lineages, showing that TL VCs have similar characteristics to those of WT SCs at the transcriptional level. **J, K**, Expression profiles of VC-required genes (**J**) and MG-required genes (**K**) in four cell lineages, showing that TL VCs have similar characteristics to those of WT SCs at the transcriptional level. Blue (**H, J**) or red (**I, K**) lines indicate the mean expression levels of all genes in the different groups. **L, M**, Gene ontology enrichment analysis of VC-required genes (**J**) and MG-required genes (**K**) in the transcriptome. ∞ indicates *P*-value=0. **N, O**, Relative expression levels of VC-preponderant or -specific genes (**N**) and MG cell-specific genes (**O**) in WT and TL-10/+ pollen. Total RNA was extracted from whole mature pollen. Results are an average of three independent experiments. Statistical significance is determined by Student’s t-test (P < 0.01, **, P < 0.001, ***) and the error bars show standard deviation (SD).

Pairwise comparisons of the transcriptomic datasets were then performed to identify differentially expressed genes (DEGs). We found that 5082 genes were significantly upregulated and 2465 genes significantly downregulated in WT SCs relative to WT VCs (WT SC vs. WT VC). Furthermore, 11962 genes were upregulated and 3932 genes downregulated in TL VCs relative to WT VCs (Figure 5, D and E). We then combined the RNA-seq datasets of WT SC vs. WT VC and TL VC vs. WT VC; this revealed 1892 VC- and 3123 MG-required genes that respectively contribute to VC or MG cell lineage specificity in male gametophytes (Figure 5, F and G). Analogous to VC- and MG-required genes, the mean expression level of the VC-peak annotated genes were downregulated and the MG-peak annotated genes were upregulated in TL VCs relative to WT VCs, which is consistent with the changes in chromatin accessibility reported above (Figures 4, I and L and 5, H-K; Supplemental Figure S6, G and H). Similar to the ATAC-seq results, gene ontology analysis revealed that VC-required genes were enriched in biological processes related to pollen tube development and signal transduction. Examples of VC-required genes include *PRK6, ANX1*, and *BUPS1*. MG-required genes, including *DUO1* and *BNB2*, were involved mainly in gene expression and organelle organization (Figure 5, L and M). Real-time quantitative PCR confirmed that *ANX1*/*2, BUPS1*/*2, RALF4*/*19, PRK1*/*2A*/*6*, and *VGD1* were downregulated, and *BNB1/2* and *DUO1* were upregulated, in TL-10/+ pollen (Figure 5, N and O). Taken together, these results are consistent with H3K27me3 erasure-induced chromatin remodeling and provide a molecular basis for the VC–MG cell fate transition suggesting that the developmental program was extensively altered in the TL VC following H3K27me3 erasure.

### TL VC exhibits MG cell ultrastructural features

Next, we analyzed the ultrastructure of TL-10 VCs to examine how sub-cellular organization was affected due to the cell fate transition caused by H3K27me3 erasure. Compared with the highly-condensed chromatin exhibited in WT SN, chromatin was highly diffuse in WT VN with no distinct nucleolus. Abnormally condensed chromatin speckles were observed in TL-10 VNs (Figure 6, C, G and I; Supplemental Figure S7). Furthermore, the lipid body, an organelle found only in WT VCs (Park and Twell, 2001), was not present in TL-10 VCs (Figure 6, B, D and H). In WT pollen, small spherical vacuoles exist in SCs, and whole vacuoles are no longer visible in VCs at the mature pollen stage (Huang et al., 2018). However, vacuoles of various sizes were present in TL VCs (Figure 6, A, B, G, H, J, O and R). The ultrastructural discrepancy between WT VC and TL VC was also observed on the simplification of organelle structure. Many regular cristae were present in the mitochondria of WT VCs; by contrast, irregular vesicular structures and shortened or absent cristae were observed in TL VCs, similar to SCs (Figure 6, E, K, M, N, S, T, W and X). In WT VCs, the Golgi body consisted of long, narrow, stacked lamellar vesicles, with smaller vesicles at each end. In TL VCs, the vesicular structures of the Golgi bodies were abnormally short and expanded. The endoplasmic reticulum had degenerated in TL VCs. These structures are analogous to that in SC (Figure 6, F, L, P, Q, U, V, Y and Z). The membrane-bound structures with fibrillar substances (MBFs) and Golgi vesicles that are considered as polysaccharide particles were absent in TL VCs, as was observed in SCs (Figure 6, D, H, M and R). Overall, there were no obvious differences in morphology or structure between WT SCs and TL SCs. The ultrastructural features of TL VCs were highly similar to those of WT and TL SCs, suggesting that the TL VCs had switched cell fate to SCs and resembled a giant version of SC.

**Figure 6.**
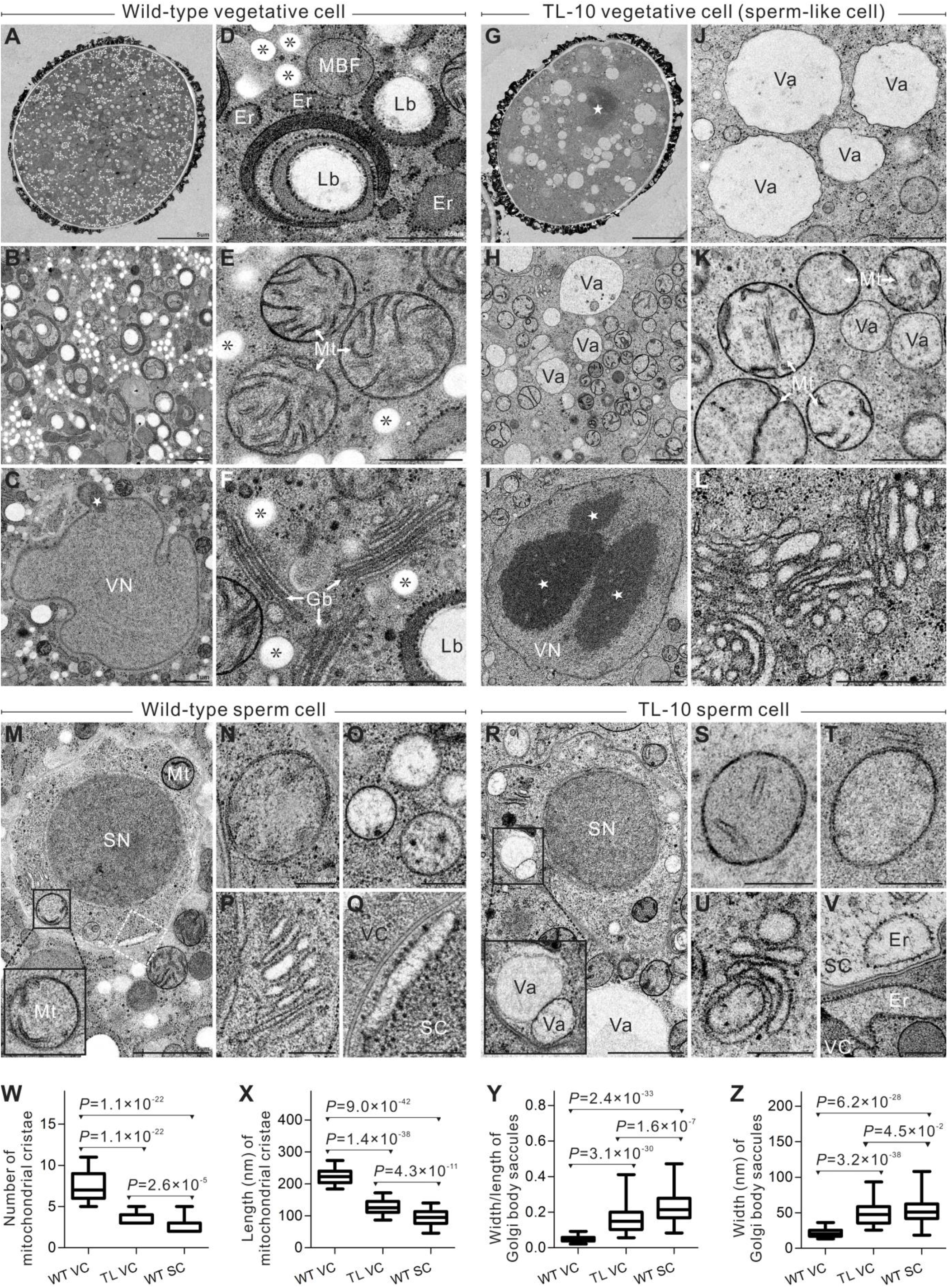
TL VCs exhibit similar ultrastructural morphology to SC. **A-F**, WT VC ultrastructure. (**A**) The panoramic image of WT VC. (**B**) Higher magnification view displays the abundant organelles in WT VCs. (**C**) Highly diffused chromatin lacking a distinct nucleolus in VN. (**D**) Golgi vesicle, membrane-bound structures with fibrillar substance, and VC-specific lipid body. (**E**) Mitochondria with numerous, regular-sized cristae. (**F**) Golgi body consisting of long, narrow, stacked lamellar vesicles with small vesicles at both ends. **G-L**, TL-10 VC ultrastructure. (**G**) The panoramic image of TL-10 VC. (**H**) Higher magnification view displays the simplification of organelles in TL-10 VCs. (**I**) The giant nucleolus and compact chromatin speckles of TL-10 VN. (**J**) Vacuoles of varying sizes. (**K**) Mitochondria with irregular vesicular structures and shortened or absent cristae. (**L**) The shortened, abnormally expanded vesicular structure in Golgi bodies. **M-V**, The ultrastructure of WT SCs (**M-Q**) and TL-10 SCs (**R-V**). (**M, R**) The panoramic images display scarce organelles and highly condensed chromatin in SCs. (**N, S, T**) The cristae are reduced or absent in SC mitochondria. (**O**) Small spherical vacuoles in SCs. (**P, U**) Golgi body in SCs. (**Q, V**) endoplasmic reticulum in SCs. (**Q**) higher magnification view of the white dotted boxes in (**M**). **W**, Number analysis of the mitochondrial cristae in WT VC, TL-10 VC and WT SC. **X**, Length analysis of mitochondrial cristae in WT VC, TL-10 VC and WT SC. The average length of the cristae is the sum of all cristae divided by the number of cristae in a mitochondrion. n=48 in WT VCs, 75 in TL VCs, and 41 in WT SCs. **Y**, Statistical analysis of length/width of Golgi body saccules in WT VC, TL-10 VC and WT SC. **Z**, Statistical analysis of width of Golgi body saccules in WT VC, TL-10 VC and WT SC. n=64 in WT VCs, 119 in TL VCs, and 83 in WT SCs. Statistical significance is determined by Student’s t-test (P < 0.05, *, P < 0.001, ***) and the error bars show standard deviation (SD). Scale bars, 5 μm (**A, G**), 1 μm (**B, C, H, I, M, R**), 0.5 μm (**D-F, J-L**), 0.2 μm (**N-Q, S-V**). SC, sperm cell; SN, sperm cell nucleus; VC, vegetative cell; VN, vegetative cell nucleus; Mt, mitochondrion; Gb, Golgi body; Er, endoplasmic reticulum; Lb, lipid body; Va, vacuole; star (☆), highly compacted chromatin speckle and nucleolus; *, Golgi vesicle.

## Discussion

The present work provided evidences supporting that H3K27me3 is essential for VC identity maintenance and H3K27me3 erasure occurred in MG cells contributes to promote SC fate establishment. By combining multi-omics approach with cytology techniques, we demonstrated that VC-targeted H3K27me3 erasure triggers the shifting of the typical VC fate to a SC fate, thus experimentally demonstrating that H3K27me3 is essential for VC fate commitment and proper pollen development. Recently, H3K27me3 erasure was reported to be critical for MG differentiation (Borg et al., 2020). Our work provides further experimental evidences for the proposal and shows its efficiency in promoting MG cell fate establishment. In fact, the differential distribution of unknown cell fate determinants has been hypothesized formerly to explain how cell fate is established in the pollen (Twell et al., 1998). Here, we reveal that H3K27me3 acts as an epigenetic determiner during pollen development to mediate cell fate distinct in VC and MG, and ensure the successful formation of the two divergent cell lineages in *Arabidopsi*s male gametophytes. An earlier study also proposed that after microspore asymmetric division, VCs maintain the ‘default setting’ of the microspore developmental program (Twell et al., 1998), but MG cells require reprograming to induce their gamete cell fate specification (Berger and Twell, 2011). Our results also reveal that H3K27me3 is one of those ‘default settings’, namely, the VC inherits the default H3K27me3 status to guarantee development and functional specification for fertilization, whereas the MG cell undergoes H3K27me3 reprogramming to determine SC fate (Figure 7).

**Figure 7.**
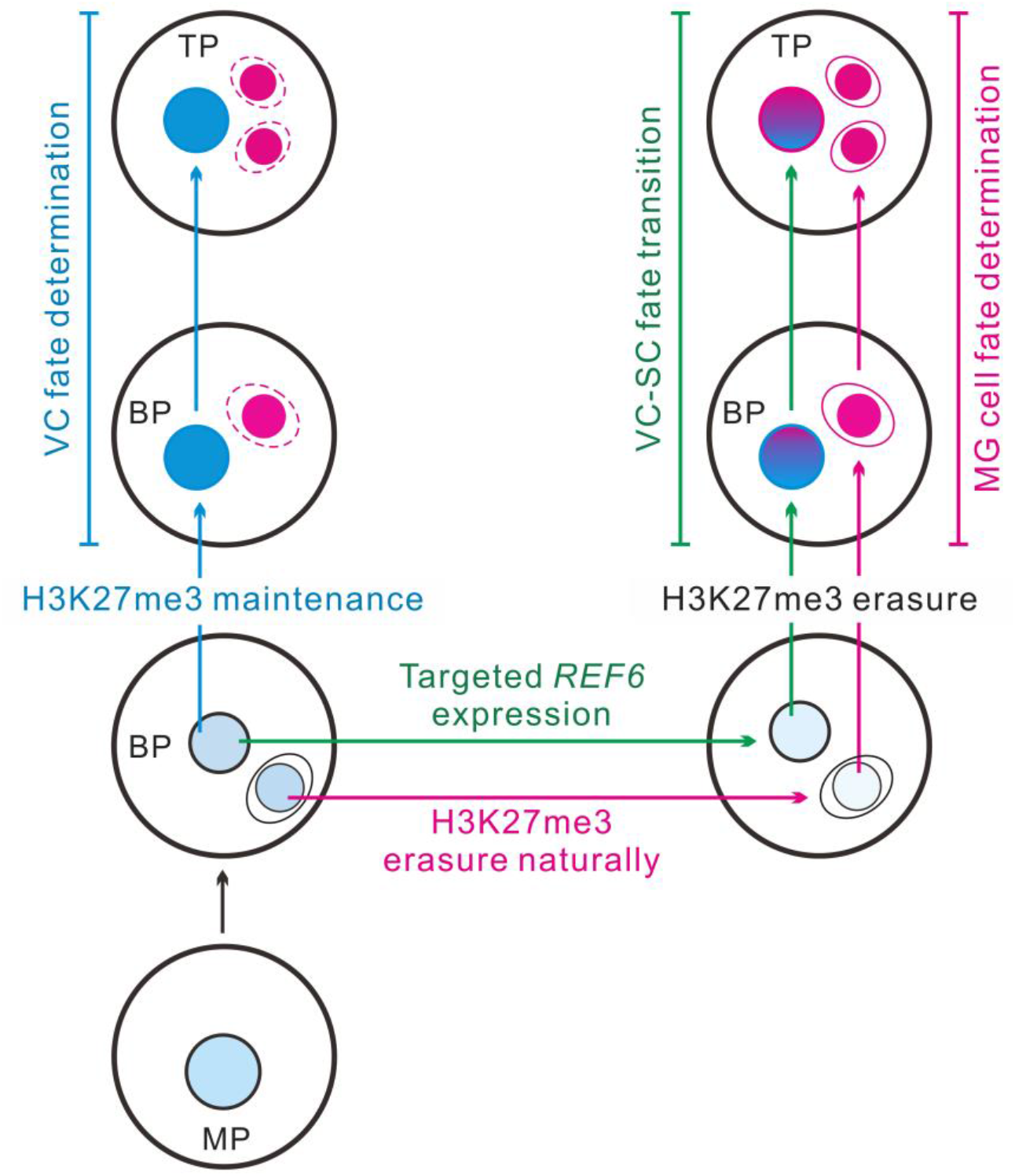
A putative model for that H3K27 methylation guides distinct cell fates of two cell lineages in male gametophyte. In angiosperm, the microspore undergoes a mitosis to generate two daughter cells. The vegetative cell (VC) intends to maintain the H3K27me3 setting from its mother cell microspore, and then differentiates into a functional male gametophyte (left, indicated in blue). On the contrary, the H3K27me3 erasure in the male germline (MG) triggers the gamete cell fate in *Arabidopsis* (right, indicated in carmine). The artificial VC-targeted H3K27me3 erasure medicated by *REF6* expression led to VC developmental reprogramming towards a MG destination, namely, the VC was shifting to SC fate (right, indicated in green). In brief, H3K27me3 acts as an epigenetic switch mediating distinct germline or vegetative cell fate in male gametophytes. To highlight VC formation pathway (left, blue), MG fate specification process in the same male gametophyte is showed by red dotted lines. MP, microspore; BP, bicellular pollen; TP, tricellular pollen.

In addition, DUO1 was the first reported regulator of GC division and MG cell fate establishment (Hackenberg and Twell, 2019). In present work, we found that *DUO1* itself was not repressed directly by H3K27me3 in VCs, however, following VC-targeted H3K27me3 removal, *DUO1* and its downstream target gene *HTR10* were ectopically expressed, and the aberrant phenotypes similar to the VC-ectopic *DUO1* expression line (*pLat52:DUO1*) (Brownfield et al., 2009) were presented in strong TL pollen grains. This suggests that despite H3K27me3 does not directly repress *DUO1* expression in VCs, it is definitely a critical upstream regulator of *DUO1*. BNB1 and BNB2, which transiently accumulate in germ cell progenitors and are required for GC specification, are the pollen-specific VIIIa subfamily bHLH transcription factors that potentially regulate *DUO1* (Yamaoka et al., 2018). Interestingly, we found that *BNB1* and *BNB2* were repressed by H3K27me3 in WT VCs and expressed in TL VCs. Thus, H3K27me3 erasure is a primary upstream signal that activates *BNB1*/*2* expression during early MG cell fate determination, which then activates *DUO1* expression.

During evolution the intricate multilayered strategies have been built up in both animals and plants to ensure that cell fate is precisely determined. In mouse, the enforced expression of a core set of transcription factors swiftly converted pluripotent stem cells into oocyte-like cells (Hamazaki et al., 2021). It is widely known that turning differentiated cells directly into another type of cells is a much harder than dedifferentiating to induce stem cells. Herein, we successfully triggered the differentiated VC shift towards a MG cell fate in one-step manipulation through VC-targeted H3K27me3 erasure and created a cell model as a useful tool to further elucidate the molecular mechanism underlying male gametogenesis, which is also an enlightenment for the researches on animal and plant reproductive biology.

As showing in present work VC-targeted H3K27me3 erasure resulted in about 15% VCs shifting towards MG cell fate, suggesting that there might be other pathways or epigenetic mechanisms regulating cell fate determination in male gametophyte. During *Arabidopsis* male gametogenesis, H3K4me3 is present in both VC and SC, but H3K9me2 and CENH3 only persist in SC (Schoft et al., 2009; Pinon et al., 2017). The SDG2-mediated H3K4me3 deposition in VCs prevents the ectopic heterochromatic H3K9me2 speckle formation and facilitates VN chromatin decondensation. The loss of H3K4me3 barely affects cell fate establishment of VCs and MG cells, but causes high compaction of chromatin in VCs and obstructs pollen germination and pollen tube elongation. Meanwhile, the H3K4me3 reduction in MG cells partly disturbs pollen mitotic cell cycle, particularly the second mitosis that sets up the two SCs (Pinon et al., 2017). These results suggest that H3K4me3 may involve in the functional specialization of VC and MG cells via a pleiotropic mechanism. The omics analysis further shows that H3K4me3 and H3K27ac accumulate selectively at the Polycomb target genes in SC, indicating that they may collaborate with H3K27me3 erasure to facilitate SC chromatin remodeling and the transcription of genes essential for male gametogenesis (Borg et al., 2020). Therefore, comprehensive epigenetic regulation network of cell fate determination in male gametophyte needs to be further explored.

## Materials and methods

### Plant material, growth conditions and transformation

All *Arabidopsis thaliana* lines are in Columbia background. Seeds were sown on 1/2 MS plates with 1.0% (w/v) sucrose and then placed in an incubator with a long-day condition (16h light/8h dark) at 22°C. About a week later, the seedlings were grown in greenhouse under a photoperiod of 16h light/8h dark at 22°C.

The HTR10-RFP marker is a gift from Prof. Frédéric Berger and *swn-21/-clf-29/+* is a gift from Prof. Xiaofeng. Cao. The *pACTIN11:H2B-GFP* reporter was obtained by the fusion of the *ACTIN11* promoter with the HISTONE2B-GFP. The DUO1-RFP marker line was constructed as described previously (Rotman et al., 2005). The *pLat52:REF6-GFP* over-expression constructs were generated via two steps. Firstly, PCR amplification of the *Lat52* promoter from the plasmid P092 (Wu et al., 2012) was performed to introduce BstXI and SacI flanking restriction sites, which was then cloned into the BstXI-SacI-digested Q108 vector (modified from vector PART27, carrying a GFP and a Hygromycin-resistance gene). Next, a wild-type genomic sequence containing *REF6* (AT3G48430) open reading frame was PCR-amplified from genomic DNA and was then cloned into the KpnI and NotI sites following the *Lat52* promoter. The gene construct was transferred into *Agrobacterium tumefaciens* strain GV3101 which was used to transform *Arabidopsis* plants by the floral dip method. For hygromycin (Merck) selection, 50μg/ml antibiotics were respectively supplemented to the media. Different independent transgenic lines were selected according to the hygromycin resistance and subsequent phenotypic analysis.

### Pollen cytological visualization

The pollen and vegetative cell morphology were conducted using a Leica SP8 confocal laser scanning microscope (CLSM, Leica Microsystems) with differential interference contrast (DIC). Mature pollen grains were dissected rapidly in Alexander’s dye (25% glycerin, 10% ethyl alcohol, 0.01% malachite green, 0.01% acid fuchsin, 0.01% Orange G, 4% acetic acid) to evaluate the substance accumulation and viability of the vegetative cell. Images were collected by an Olympus IX71 microscope. Unless otherwise indicated in the figure legends, the images come from one representative of at least three independent experiments. For every experiment, pollen grains from at least five flowers of the control and TL lines were collected and stained.

### Pollen germination assays

The *in vitro* pollen germination was performed according to previous method (Wu et al., 2012). The pollen grains were harvested from fresh self-dehiscing anthers and placed onto pollen germination medium (PGM: 18% sucrose, 1mM CaCl_2_·2H_2_O, 1mM Ca(NO_3_)_2_·4H_2_O, 1mM MgSO_4_·7H_2_O, 1% H_3_BO_3_, pH7.0, containing 0.9% agarose) and cultured at 22°C for 4h. For *in vivo-in vitro* pollen growth assay, wild-type styles were cut horizontally at the shoulder region of the ovaries after hand-pollination by control and TL pollen grains respectively. Then, the pollinated styles were incubated on PGM for 6h. Unless otherwise indicated in figure legends, the images are from one representative of at least three independent experiments. It is worth emphasizing that the materials from control and different TL lines were placed in the same culture dish for every experiment to guarantee the consistency of experimental conditions.

### Cryo-fixation, freeze-substitution and transmission electron microscopy (TEM)

The dehiscing anthers encased with mature pollen grains were dissected into the high-pressure freezer specimen holder and then embedded with the external cryo-protectant (2% low melting point agarose, Sigma, A9539). During the manipulation any air bubbles should be carefully removed since they would collapse under high-pressure and cause disruptions to the tissues. Samples were rapidly freeze fixed using the Lene and stoica EM ICE and stored in liquid nitrogen.

Fixative can be conveniently prepared. We made stock solutions of 4% osmium tetroxide (OsO_4_) in acetone then stored them in liquid nitrogen. OsO_4_ crystals are dissolved in a special glass container. We put three specimen holders carrying samples (three biological repeats) into one standard 1.8 ml cryo-tubes for the freeze substitution. Samples at liquid nitrogen temperature were transferred to fixative vials at identical temperature, and then placed in the Leica EM ASF2 where they were warmed to −78.5°C. Substitution was completed in 2.5 day, and then the samples were warmed according to the following schedule: 2 hr at −20°C (standard freezer), 2 hr at 0-4°C (refrigerator), and 2 hr at 23°C (room temperature). Samples were separated charily from the holders and rinsed three times in dried acetone. After epoxy propane gradient replacement, samples were embedded in Spurr resin. Ultra-thin sections preparation and subsequent TEM observation was performed following a previous description (Huang et al., 2019). Ultra-thin sections (70 nm) were stained with 2% (w/v) uranyl acetate and 2.6% (w/v) lead citrate aqueous solution and then examined with a JEM-1400 plus transmission electron microscope (JEOL) at 100 kV.

### Pollen grains collection

Fresh flowers were cut and placed in a 50 ml centrifuge tube containing 25ml pollen buffer (PB: 18% sucrose, 1% H_3_BO_3_, 1mM CaCl_2_·2H_2_O, 1mM Ca(NO_3_)_2_·4H_2_O, 1mM MgSO_4_·7H_2_O, pH7.0). Oscillate violently for 5-10 min to ensure that the pollen grains could be washed out. Filter the pollen suspension through a 75μm nylon mesh and then pellet pollen grains at room temperature (RT), 1500rpm, 3min.

### Quantitative real-time PCR assay

Total RNA of mature pollen grains was extracted using the TaKaRa MiniBEST Plant RNA Extraction Kit (9769) according to the manufacturer’s protocol. The subsequent process was performed following a previous description (Huang et al., 2018; Huang et al., 2019). The first strand of cDNA was synthesized using PrimeScript™RT Master Mix (Perfect Real Time) (TaKaRa). Quantitative real-time PCR (RT-qPCR) assay was performed using FastStart Essential DNA Green Master (Roche) on a CFX Connect™ Real-Time System (Bio-Rad). Each experiment was repeated three times and samples were normalized according to the *UBQ10* expression. Data acquisition used Bio-Rad CFX Manager software; the relative expression levels were measured by the 2^(–ΔΔCt)^ analysis method (Huang et al., 2018; Huang et al., 2019).

### Whole-mount immunofluorescence

Whole-mount immunofluorescence of histone modifications in the pollen grains was adapted from a procedure described previously (She et al., 2018). Pollen grains were immediately fixed with 1% formaldehyde containing 0.1% Triton X-100 for 30 min with gentle rotation, and then rinsed two times in 1×TTBS buffer (TBS, 0.1% Triton X-100). Before antibody labeling samples pretreatment included tissue clarification [ethanol-xylene (1:1), RT, 30 min; rinsed 2×10 min in 1×TTBS buffer], cell wall digestion (1.5% cellulose R-10, 0.75% macerozyme R-10, 28°C, 4h) and blocking (TBS, 0.1% Triton X-100, 5% BSA, RT, 1h) were performed. Following, samples were incubated with primary antibodies (anti-H3K27me3, Millipore 07-449, 1:100; and anti-H3K9me2, Abcam ab1220, 1:100) for 24h at 4°C and washed in 1×TTBS, two times. Then samples were incubated with either anti-rabbit Alexa Fluor 647 or 488 (Invitrogen, for anti-H3K27me3), or anti-mouse Alexa Fluor 647 (Invitrogen, for anti-H3K9me2) for 24h at 4°C and washed in 1×TTBS, two times. Finally, pollen grains were spotted onto slides and mounted in Vectashield mounting medium with DAPI (Vector Laboratories). Images were taken using a Leica SP8 CLSM.

### Nuclei isolation and fluorescence-activated nuclei sorting (FANS)

TL-10/+ carrying the HTR10-RFP marker and control (*pHTR10:HTR10-RFP*; *pACTIN11:H2B-GFP* double-labeled line) carry GFP and RFP fluorescent markers in the SN and VN respectively, thus sorting of viable SN and VN is efficient. The collected pollen grains were re-suspended in Galbraith buffer (30mM Trisodium citrate dehydrate, 45mM MgCl_2_, 20mM MOPS, 0.1% (v/v) Triton X-100, adjust pH to 7.0) and then transferred to a 1.5 ml tube containing glass beads, before vortexed continuously at 2000rpm for 3 min. Filter the sample through a 10μm nylon mesh in a 2ml tube to acquire the nucleus mixture.

Fluorescence-activated SN and VN sorting were performed in a BD FACSAria III cell sorter. The nuclei can be obtained with high purity and integrity by running a 100μm nozzle with 330kPa sheath pressure. Use GFP (488nm) and RFP (561nm) to gate VN population (GFP-positive) and SN population (RFP-positive) respectively. Gate also for both VN and SN with a uniform side scatter (SSC) profile, which allowed the exclusion of debris and incomplete nuclei. The sorted nuclei were collected directly into Galbraith buffer and kept on ice for the subsequent experiments. The FANS plot analysis was performed by FACSDiva software (BD Biosciences).

### Microspore isolation and fluorescence-activated cell sorting (FACS)

Flower buds at late microspore stage were picked from TL-10/+ carrying the HTR10-RFP marker and control (*pHTR10:HTR10-RFP line*) plants. The sample preparation and FACS of the late microspore (LMSs) on a BD FACSAria III cell sorter were preformed according to the method described previously (Santos et al., 2017) with minor modifications. The sorter was run in standard configuration with a 100μm nozzle, and gate for microspores based on their green autofluorescence (488nm). Microspores also gate with the RFP (561nm) channel to allow the exclusion of unwished bicellular pollen. The collected microspores were maintained in ice and RNA was extracted immediately.

### Cleavage under targets and tagmentation (CUT&Tag) and data analysis

CUT&Tag assay was employed to profiling H3K27me3 as described previously with minor modifications (Kaya-Okur et al., 2019). The FANS-purified nuclei must be preserved in ice to maintain fresh and good viability, but never frozen. About 10^6^ nuclei per sample was collected and then centrifuged at 500 g at 4°C for 10 min. Aliquots of nuclei were washed twice in 0.5 ml Wash Buffer (20mM HEPES pH 7.5; 150mM NaCl; 0.5mM Spermidine; 1× Protease inhibitor cocktail) by gentle pipetting, then centrifuged for removing the supernatant. Concanavalin A-coated magnetic beads (Bangs Laboratories) were washed twice with binding buffer (20 mM HEPES pH 7.5, 10 mM KCl, 1 mM MnCl_2_, 1 mM CaCl_2_) for activation, and then 10 μl beads were added per sample and subsequently incubated at RT for 15 min. Bead-bound cells were re-suspended in 50 μl Dig-wash Buffer (20mM HEPES pH 7.5; 150mM NaCl; 0.5mM Spermidine; 1× Protease inhibitor cocktail; 0.05% Digitonin) containing 2mM EDTA and a 1:50 dilution of primary antibody (anti-H3K27me3, Millipore), subsequently incubated overnight at 4°C with slow rotation. Removed primary antibody thoroughly and added secondary antibody (Santa Cruz) diluted in 50μl Dig-wash Buffer (1:100), and then incubated at RT for 1 h. Then nuclei were washed using magnet stand 3× for 5 min in 1ml Dg-Wash buffer to remove the unbound antibodies. A 1:100 dilution of pA-Tn5 adapter complex (TTE mix, 4μM, Novoprotein) was prepared in Dig-med Buffer (20mM HEPES, pH 7.5, 300mM NaCl, 0.5mM Spermidine, 1× Protease inhibitor cocktail, 0.05% Digitonin). Nuclei were incubated with 100μl 0.04 μM TTE mix at RT for 1h with gentle vortexing, and then washed 3× for 5 min in 1ml Dig-med Buffer to remove the unbound pA-Tn5 protein. The nuclei were then re-suspended in 300μl Tagmentation buffer (10mM MgCl_2_ in Dig-med Buffer) and incubated at 37°C for 1 h. To terminate tagmentation, 10μl 0.5M EDTA, 2.5μl 20mg/ml Proteinase K and 3μl 10% SDS were added to samples and incubated at 55°C for 1h. DNA was purified using phenol-chloroform-isoamyl extraction and ethanol precipitation, as well as RNaseA treatment.

For library amplification, 21 μl DNA was mixed with 2 μl of a universal i5 and a uniquely barcoded i7 primer (Buenrostro et al., 2015), and then 25 μl NEB Next HiFi 2× PCR Master mix was added and mixed. A total volume of 50μl sample was placed in a PCR thermal cycler with a heated lid via the following program: 72°C for 5min; 98°C for 30s; 16 cycles of 98°C for 10s and 63°C for 30s; final extension at 72°C for 1 min and hold at 4°C. To purify PCR products, 1.5 × volumes of AMPure XP beads (Beckman Coulter) were added and incubated for 15 min at RT. Next, the libraries were washed twice gently in 80% ethanol, and eluted in 30μl 10mM Tris pH 8.0. The size distribution of libraries was determined by Agilent 4200 TapeStation analysis, and all libraries were performed Paired-end Illumina sequencing on an Illumina NovaSeq platform.

Before mapping of pair-end reads, clean reads were obtained from the raw reads by removing the adaptor sequences and then aligned to the *Arabidopsis* genome reference sequences (TAIR10) using the BWA program. We calculate the fragment sizes for read pairs given a BAM file from paired-end sequencing. Several regions were sampled to estimate the summary statistics on the fragment lengths. Properly paired reads were used for computation. The BAM file generated by the unique mapped reads as an input file, using MACS2 software for callpeak with cutoff q-value < 0.05 and the parameters “macs2 callpeak--nomodel-f BAMPE--keep-dup 1-q 0.05 -B-SPMR” (Zhang et al., 2008). Reads distributions from bigwig across peaks and genes were presented as an average plot (average signals of reads across all peaks and genes) using the deeptools tool. The peaks annotation and genomic distribution of CUT&Tag peaks were analyzed by HOMER (Heinz et al., 2010), and we counted the annotations results and plotted the distribution results by R software. Independent biological replicates showed similarity by Pearson correlation coefficient analysis. CUT&Tag signal tracks were presented by Integrative Genomics Viewer (IGV) software (Robinson et al., 2011).

### Assay for transposase accessible chromatin (ATAC)-seq and data analysis

ATAC-seq is a method for mapping chromatin accessibility genome-wide and used to conclude the regions of increased accessibility. Transposase integration reactions and sequencing library preparations were manipulated as a procedure reported previously (Buenrostro et al., 2015). It is worth of emphasizing that FACS-purified fresh nuclei to be used for ATAC-seq must be placed in ice prior to transposase integration reaction but kept never frozen. Approximately 10^6^ nuclei per sample was collected and pelleted by centrifugation at 500 g at 4°C for 10 min. The supernatant was then removed, leaving ∼50 μl in the tube. The nuclei were re-suspended and washed with 1× phosphate buffered saline (PBS), centrifuged using the above setting, and the supernatant removed. Next, the Tn5 reaction was prepared and mixed well with nuclei using the Nextera reagents (Illumina, FC-121-1030) as follows: 25 μl 2 × TD buffer, 2.5 μl TDE1 (Nextera Tn5 Transposase) and nuclease-free H_2_O were added to total 50 μl. The reaction was placed in a thermal cycler at 37°C for 30 min. DNA fragments were purified immediately using the Minelute PCR purification kit (Qiagen) and eluted in 10 μl elution buffer which consists of 10 mM Tris-HCl (pH 8), and each sample underwent amplified using NEB Next High-Fidelity 2 × PCR Master Mix for 12 PCR cycles. All amplified ATAC-seq libraries were purified using AMPure XP beads (Beckman Coulter), quantified by a Qubit Fluorometer and analyzed on an Agilent 2100 Bioanalyzer prior to pooling and subsequently sequenced using Illumina NovaSeq.

Mapping of pair-end reads, callpeak, peaks annotation and genomic distribution of ATAC-seq peaks were analyzed as described in CUT&Tag analysis. The principal component analysis (PCA) was performed on the signals of merged peaks from all samples, and the PlotPCA sorted the principal components according to the variability of the data. Based on this, a plot which shows the eigenvalues of the top two principal components was obtained. Merge the peak files of all replicate samples from each cell type was using the Bedtools software and the differential accessible regions (DARs) were assessed using DESeq2 (Love et al., 2014) according to absolute value of log_2_ fold change (FC) was 1 (|log_2_FC| > 1) at P-value < 0.05. DARs were annotated by the function of annotatePeak of ChIPseeker (Yu et al., 2015). The distributions of the reads across genes were presented using the deeptools based on the BED file.

### RNA-seq and data analysis

Messenger RNA (mRNA) isolation and RNA-Seq library construction was performed as the procedure described previously (Zhao et al., 2019). mRNA was extracted from the FANS-isolated VC nuclei and SC nuclei of control and TL-10/+ (about 10^6^ nuclei per sample) with independent biological replicates using a Dynabeads™ mRNA DIRECT™ Micro Purification Kit (Life Technologies). Total RNA of late microspore (about 10^6^ microspores per sample) was extracted using a RNeasy Plant Mini Kit (QIAGEN). Subsequent cDNA synthesis and amplification were performed using a SMART-Seq® HT Kit (Takara), and then purified using an Agencourt AMPure Purification Kit (Beckman Coulter). Purified cDNA was quantified by a Qubit 3.0 Fluorometer (Life Technologies) and validated using a Bioanalyzer 2100 system (Agilent Technologies). Sequencing libraries were constructed using the VAHTS Universal V6 RNA-seq Library Kit for MGI (Vazyme) following the manufacturer’s recommendations with unique index codes. The Library quantification and size was assessed using a Qubit 3.0 Fluorometer and Bioanalyzer 2100 system. Subsequently, sequencing was performed on MGI-SEQ 2000 platform by Frasergen Bioinformatics Co., Ltd. (Wuhan, China) and obtained about 6 Gb raw sequence data per library.

Low quality reads, such as reads with adaptor sequences, reads with >5% N, or >20% bases and with quality <Q20 (percentage of sequences with sequencing error rates <1%), were removed using perl script. The clean reads filtered from raw reads were mapped to the *Arabidopsis* genome (TAIR10) using HISAT2 (Li and Dewey, 2011), and gene expression levels were quantified and normalized as fragments per kilo-base per million bases (FPKM) measurements using RSEM (Kim et al., 2019). RNA-Seq heatmaps were generated using pheatmap package in R (https://github.com/raivokolde/pheatmap). DESeq2 (Love et al., 2014) was used to identify differentially expressed genes (DEGs) between sample groups, and only the genes with a fold change value≥1 (|log_2_FC|≥1) and false discovery rate (FDR) significance score (padj) < 0.01 were considered as significant DEGs. Hierarchical clustering of the gene expression data was carried out using the Agnes function in the Cluster package based on the value of the Pearson correlation coefficient between different samples.

### Gene ontology (GO) analysis

GO analysis was performed to facilitate elucidating the biological implications of the unique genes (Ashburner et al., 2000). We downloaded the GO terms for the genes from The Arabidopsis Information Resource (TAIR) (https://www.arabidopsis.org/). Fisher’s exact test (ATAC-seq) or Hypergeometric distribution (RNA-seq) was applied to identify the GO categories and false discovery rate (FDR) method was used to correct the P-values.

## Data availability statement

High-throughput sequencing data that support the findings of this study have been available in the NCBI Gene Expression Omnibus under accession code GSE162640.

## Supplemental data

The following materials are available in the online version of this article.

**Supplemental Figure 1** Comparison of male gametophyte development and function between wild-type (WT) and clf-29/+ swn-21/-double mutant.

**Supplemental Figure 2** TL line selection and H3K27me3 is essential for VC development.

**Supplemental Figure 3** H3K27me3 is indispensable for VC functional specification.

**Supplemental Figure 4** H3K27me3 CUT & Tag profiling of Arabidopsis WT VC, WT SC and TL VC.

**Supplemental Figure 5** ATAC-seq profiling of Arabidopsis WT VC, WT SC, TL VC and TL SC and global dynamics of chromatin accessibility during VC and MG lineage development.

**Supplemental Figure 6** Transcriptome reproducibility, correlation among cell lineages and transcriptome dynamics during VC and MG lineage development.

**Supplemental Figure 7** Abnormal condensation of chromatin in TL-10 VN.

## Acknowledgements

We thank Prof. Xiaofeng Cao (Institute of Genetics and Developmental Biology, Chinese Academy of Sciences) for the *swn-21/-clf-29/+* seed; Prof. Frédéric Berger (Gregor Mendel Institute, Austrian Academy of Sciences) for the HTR10-RFP marker seed and careful reading of the manuscript; Dr. Wenxuan Zou (College of Life Science, Wuhan University) for Cryo-fixation, freeze-substitution and TEM; Dr. Ying Liu (Medical Research Institute, Wuhan University) for FANS; Shanghai Jiayin Biotechnology for H3K27me3 CUT&Tag and ATAC-seq library construction, sequencing and data analysis; Wuhan Frasergen Bioinformatics Co., Ltd. for RNA-Seq library construction, sequencing and data analysis.

## Parsed Citations

Ashburner M, Ball CA, Blake JA, Botstein D, Butler H, Cherry JM, Davis AP, Dolinski K, Dwight SS, Eppig JT, Harris MA, Hill DP, Issel-Tarver L, Kasarskis A, Lewis S, Matese JC, Richardson JE, Ringwald M, Rubin GM, Sherlock G (2000) Gene ontology: tool for the unification of biology. The Gene Ontology Consortium. Nat Genet 25(1): 25–29.

Bayer M, Nawy T, Giglione C, Galli M, Meinnel T, Lukowitz W (2009) Paternal control of embryonic patterning in Arabidopsis thaliana. Science 323(5920): 1485–1488.

Berger F, Twell D. (2011) Germline specification and function in plants. Annu Rev Plant Biol 62: 461–84.

Borg M, Brownfield L, Khatab H, Sidorova A, Lingaya M, Twell D (2011) The R2R3 MYB transcription factor DUO1 activates a male germline-specific regulon essential for sperm cell differentiation in Arabidopsis. Plant Cell 23: 534–549.

Borg M, Berger F (2015) Chromatin remodeling during male gametophyte development. Plant J 83(1): 177–188.

Borg M, Jacob Y, Susaki D, LeBlanc C, Buendía D, Axelsson E, Kawashima T, Voigt P, Boavida L, Becker J, Higashiyama T, Martienssen R, Berger F (2020) Targeted reprogramming of H3K27me3 resets epigenetic memory in plant paternal chromatin. Nat Cell Biol 22(6): 621–629.

Boisson-Dernier A, Roy S, Kritsas K, Grobei MA, Jaciubek M, Schroeder JI Grossniklaus U (2009) Disruption of the pollen-expressed FERONIAhomologs ANXUR1 and ANXUR2 triggers pollen tube discharge. Development 136(19): 3279–3288.

Brownfield L, Hafidh S, Borg M, Sidorova A, Mori T, Twell D (2009) Aplant germline-specific integrator of sperm specification and cell cycle progression. PLoS Genet 5: e1000430.

Buenrostro JD, Wu B, Litzenburger UM, Ruff D, Gonzales ML, Snyder MP, Chang HY, Greenleaf WJ (2015) Single-cell chromatin accessibility reveals principles of regulatory variation. Nature 523(7561): 486–90.

Buenrostro JD, Wu B, Chang HY, Greenleaf WJ (2015) ATAC-seq: A method for assaying chromatin accessibility genome-wide. Curr Protoc Mol Biol 5(109): 21.29.1–21.29.9.

Chen Z, Hafidh S, Poh SH, Twell D, Berger F (2009) Proliferation and cell fate establishment during Arabidopsis male gametogenesis depends on the retinoblastoma protein. Proc Natl Acad Sci USA 106: 7257–7262.

Chen Z, Jeanie LH, Mathieu I, Venkatesan S, Berger F (2008) Chromatin assembly factor 1 regulates the cell cycle but not cell fate during male gametogenesis in Arabidopsis thaliana. Development 135: 65–73.

Calarco JP, Borges F, Donoghue MT, Van Ex F, Jullien PE, Lopes T, Gardner R, Berger F, Feijó JA, Becker JD, Martienssen RA (2012) Reprogramming of DNA methylation in pollen guides epigenetic inheritance via small RNA. Cell 151: 194–205.

Cui X, Lu F, Qiu Q, Zhou B, Gu L, Zhang S, Kang Y, Cui X, Ma X, Yao Q, Ma J, Zhang X, Cao X (2016) REF6 recognizes a specific DNA sequence to demethylate H3K27me3 and regulate organ boundary formation in Arabidopsis. Nat Genet 48(6): 694–699.

Ge Z, Bergonci T, Zhao Y, Zou Y, Du S, Liu MC, Luo X, Ruan H, García-Valencia LE, Zhong S, Hou S, Huang Q, Lai L, Moura DS, Gu H, Dong J, Wu HM, Dresselhaus T, Xiao J, Cheung AY, Qu LJ (2017) Arabidopsis pollen tube integrity and sperm release are regulated by RALF-mediated signaling. Science 358(6370): 1596–1600.

Hackenberg D, Twell D. (2019) The evolution and patterning of male gametophyte development. Curr Top Dev Biol 131: 257–298.

Hamazaki N, Kyogoku H, Araki H, Miura F, Horikawa C, Hamada N, Shimamoto S, Hikabe O, Nakashima K, Kitajima TS, Ito T, Leitch HG, Hayashi K (2021) Reconstitution of the oocyte transcriptional network with transcription factors. Nature 589(7841):264-269.

Heinz S, Benner C, Spann N, Bertolino E, Lin YC, Laslo P, Cheng JX, Murre C, Singh H, Glass CK (2010) Simple combinations of lineage determining transcription factors prime cis-regulatory elements required for macrophage and B cell identities. Mol Cell 38: 576–589.

Higo A, Kawashima T, Borg M, Zhao M, López-Vidriero I, Sakayama H, Montgomery SA, Sekimoto H, Hackenberg D, Shimamura M, Nishiyama T, Sakakibara K, Tomita Y, Togawa T, Kunimoto K, Osakabe A, Suzuki Y, Yamato KT, Ishizaki K, Nishihama R, Kohchi T, Franco-Zorrilla JM, Twell D, Berger F, Araki T (2018) Transcription factor DUO1 generated by neo-functionalization is associated with evolution of sperm differentiation in plants. Nat Commun 9(1): 5283.

Huang X, Peng X, Xie F, Mao W, Chen H, Sun, M-X (2018) The stereotyped positioning of the generative cell associated with vacuole dynamics is not required for male gametogenesis in rice pollen. New Phytol 218(2): 463–469.

Huang X, Run M, Sun M-X (2019) OsGCD1, a novel player in rice intine construction. J Genet Genomics 46(7): 359–362.

Ingouff M, Rademacher S, Holec S, Soljić L, Xin N, Readshaw A, Foo SH, Lahouze B, Sprunck S, Berger F (2010) Zygotic resetting of the HISTONE 3 variant repertoire participates in epigenetic reprogramming in Arabidopsis. Curr Biol 20: 2137–2143.

Jackson JP, Lindroth AM, Cao X, Jacobsen SE (2002) Control of CpNpG DNA methylation by the KRYPTONITE histone H3 methyltransferase. Nature 416(6880): 556–560.

Jiang L, Yang SL, Xie LF, Puah CS, Zhang XQ, Yang WC, Sundaresan V, Ye D (2005) VANGUARD1 encodes a pectin methylesterase that enhances pollen tube growth in the Arabidopsis style and transmitting tract. Plant Cell 17(2): 584–596.

Johnson L, Cao X, Jacobsen S (2002) Interplay between two epigenetic marks. DNA methylation and histone H3 lysine 9 methylation. Curr Biol 12(16): 1360–1367.

Kaya-Okur HS, Wu SJ, Codomo CA, Pledger ES, Bryson TD, Henikoff JG, Ahmad K, Henikoff S (2019) CUT&Tag for efficient epigenomic profiling of small samples and single cells. Nat Commun 10: 1930.

Kim D, Paggi JM, Park C, Bennett C, Salzberg SL (2019) Graph-based genome alignment and genotyping with HISAT2 and HISAT-genotype. Nat Biotechnol 37(8): 907–915.

Li B, Dewey CN (2011) RSEM: accurate transcript quantification from RNA-Seq data with or without a reference genome. BMC Bioinformatics 12: 323.

Liu C, Lu F, Cui X, Cao X (2010) Histone methylation in higher plants. Annu Rev Plant Biol 61: 395e420.

Love MI, Huber W, Anders S (2014) Moderated estimation of fold change and dispersion for RNA-seq data with DESeq2. Genome Biol 15: 550.

Lu F, Cui X, Zhang S, Jenuwein T, Cao X (2011) Arabidopsis REF6 is a histone H3 lysine 27 demethylase. Nat Cenet 43(7): 715–719.

Park SK, Twell D (2001) Novel patterns of ectopic cell plate growth and lipid body distribution in the Arabidopsis gemini pollen1 mutant. Plant Physiol 126(2): 899–909.

Pinon V, Yao X, Dong A, Shen WH (2017) SDG2-mediated H3K4me3 is crucial for chromatin condensation and mitotic division during male gametogenesis in Arabidopsis. Plant Physiol 174(2): 1205–1215.

Reik W, Dean W, Walter J (2001) Epigenetic reprogramming in mammalian development. Science 293: 1089–1093.

Robinson JT, Thoryaldsdóttir H, Winckler W, Guttman M, Lander ES, Getz G, Mesirov JP (2011) Integrative genomics viewer. Nat Biotechnol 29: 24–26.

Rotman N, Durbarry A, Wardle A, Yang WC, Chaboud A, Faure JE, Berger F, Twell D (2005) Anovel class of MYB factors controls sperm cell formation in plants. Curr Biol 15: 244–248.

Santos MR, Bispo C, Becker JD (2017) Isolation of Arabidopsis Pollen, Sperm Cells, and Vegetative Nuclei by Fluorescence-Activated Cell Sorting (FACS). Methods Mol Biol 1669: 193–210.

Sassone-Corsi P (2002) Unique chromatin remodeling and transcriptional regulation in spermatogenesis. Science 296(5576): 2176–2178.

Schoft VK, Chumak N, Mosiolek M, Slusarz L, Komnenovic V, Brownfield L, Twell D, Kakutani T, Tamaru H (2009) Induction of RNA directed DNA methylation upon decondensation of constitutive heterochromatin. EMBO Rep 10(9): 1015–1021.

She W, Baroux C, Grossniklaus U (2018) Cell-type specific chromatin analysis in whole-mount plant tissues by immunostaining. Methods Mol Biol 1675: 443–454.

Takeuchi H, Higashiyama T (2016) Tip-localized receptors control pollen tube growth and LURE sensing in Arabidopsis. Nature 531: 245–248.

Twell D, Park SK, Lalanne E (1998) Asymmetric division and cell-fate determination in developing pollen. Trends Plant Sci 3: 305–310.

Twell D, Park SK, Hawkins TJ, Schubert D, Schmidt R, Smertenko A, Hussey, PJ (2002) MOR1/GEM1 has an essential role in the plant-specific cytokinetic phragmoplast. Nat Cell Biol 4(9): 711–714.

Wu JJ, Peng XB, Li WW, He R, Xin HP, Sun M-X (2012) Mitochondrial GCD1 dysfunction reveals reciprocal cell-to-cell signaling during the maturation of Arabidopsis female gametes. Dev Cell 23: 1–16.

Yamaoka S, Nishihama R, Yoshitake Y, Ishida S, Inoue K, Saito M, Okahashi K, Bao H, Nishida H, Yamaguchi K, Shigenobu S, Ishizaki K, Yamato KT, Kohchi T (2018) Generative cell specification requires transcription factors evolutionarily conserved in land plants. Curr Biol 8: 479–486.

Yu GC, Wang LG, He Q-Y (2015) ChIPseeker: an R/Bioconductor package for ChIP peak annotation, comparison and visualization. Bioinformatics 31(14): 2382–2383.

Zhang Y, Liu T, Meyer CA, Eeckhoute J, Johnson DS, Bernstein BE, Nusbaum C, Myers RM, Brown M, Li W, Liu XS (2008) Model-based analysis of ChIP-seq (MACS). Genome Biol 9: R137.

Zhao P, Zhou XM, Shen K, Liu ZZ, Cheng TH, Liu DN, Cheng YB, Peng XB, Sun M-X (2019) Two-step maternal-to-zygotic transition with two-phase parental genome contributions. Dev Cell 49: 1–12.

